# Variability in human drug metabolizing cytochrome P450 CYP2C9, CYP2C19 and CYP3A5 activities caused by genetic variations in cytochrome P450 oxidoreductase

**DOI:** 10.1101/640540

**Authors:** Maria Natalia Rojas Velazquez, Shaheena Parween, Sameer S Udhane, Amit V Pandey

**Author notes:** Address for correspondence: PD Dr. Amit V Pandey, KIKL C837, Pediatric Endocrinology, Diabetology & Metabolism, University Children’s Hospital Bern, Freiburgstrasse 15, CH-3010 Bern Switzerland. Web: http://www.pandeylab.org. These authors contributed equally to the manuscript (Maria Natalia Rojas Velazquez, Shaheena Parween). Current address: Department of Pathology, Medical College of Wisconsin Cancer Center, Milwaukee, WI 53226, USA.

## Abstract

A broad spectrum of human diseases are caused by mutations in the NADPH cytochrome P450 oxidoreductase (POR). Cytochrome P450 proteins perform several reactions, including the metabolism of steroids, drugs, and other xenobiotics. In 2004 the first human patients with defects in POR were reported, and over 250 variations in POR are known. Information about the effects of POR variants on drug metabolizing enzymes is limited and has not received much attention. By analyzing the POR sequences from genomics databases, we identified potentially disease-causing variations and characterized these by *in vitro* functional studies using recombinant proteins. Proteins were expressed in bacteria and purified for activity assays. Activities of cytochrome P450 enzymes were tested *in vitro* using liposomes prepared with lipids into which P450 and P450 reductase proteins were embedded. Here we are reporting the effect of POR variants on drug metabolizing enzymes CYP2C9, CYP2C19, and CYP3A5 which are responsible for the metabolism of many drugs. POR Variants A115V, T142A, A281T, P284L, A287P, and Y607C inhibited activities of all P450 proteins tested. Interestingly, the POR variant Q153R showed a reduction of 20-50% activities with CYP2C9 and CYP2C19 but had a 400% increased activity with CYP3A5. The A287P is most common POR mutation found in patients of European origin, and significantly inhibited drug metabolism activities which has important consequences for monitoring and treatment of patients. *In vitro,* functional assays using recombinant proteins provide a useful model for establishing the metabolic effect of genetic mutations. Our results indicate that detailed knowledge about POR variants is necessary for correct diagnosis and treatment options for persons with POR deficiency and the role of changes in drug metabolism and toxicology due to variations in POR needs to be addressed.

## Introduction

Cytochromes P450 proteins belong to heme-thiolate monooxygenase family and catalyze the conversion of many drugs and xenobiotics into the corresponding hydroxylated metabolites [1]. There are two types of cytochrome P450 in humans, type I (located in mitochondria), which participate in the biosynthesis of steroids, sterols, and retinoids and rely on ferredoxin reductase and ferredoxin as their redox partners [2,3]. The type 2 cytochromes P450 (located in the endoplasmic reticulum) participate in the metabolism of drugs and xenobiotics and rely on reduced nicotinamide adenine dinucleotide phosphate (NADPH) for the supply of redox equivalents via NADPH cytochrome P450 oxidoreductase (POR) [4,5,6]. POR is a flavoprotein which is essential in drug metabolism due to its role as an obligatory redox partner for all cytochrome P450 proteins located in the endoplasmic reticulum [4]. POR has flavin adenine dinucleotide (FAD) and flavin mononucleotide (FMN) domains that are connected by a flexible hinge region (residues 235 to 247) and electrons are transferred from NADPH through FAD hydroquinone to the isoalloxazine ring of FMN which then transfers it one at a time to the redox partners like cytochromes P450 [4,7,8,9,10,11] (**Figure 1**).

**Figure 1:**
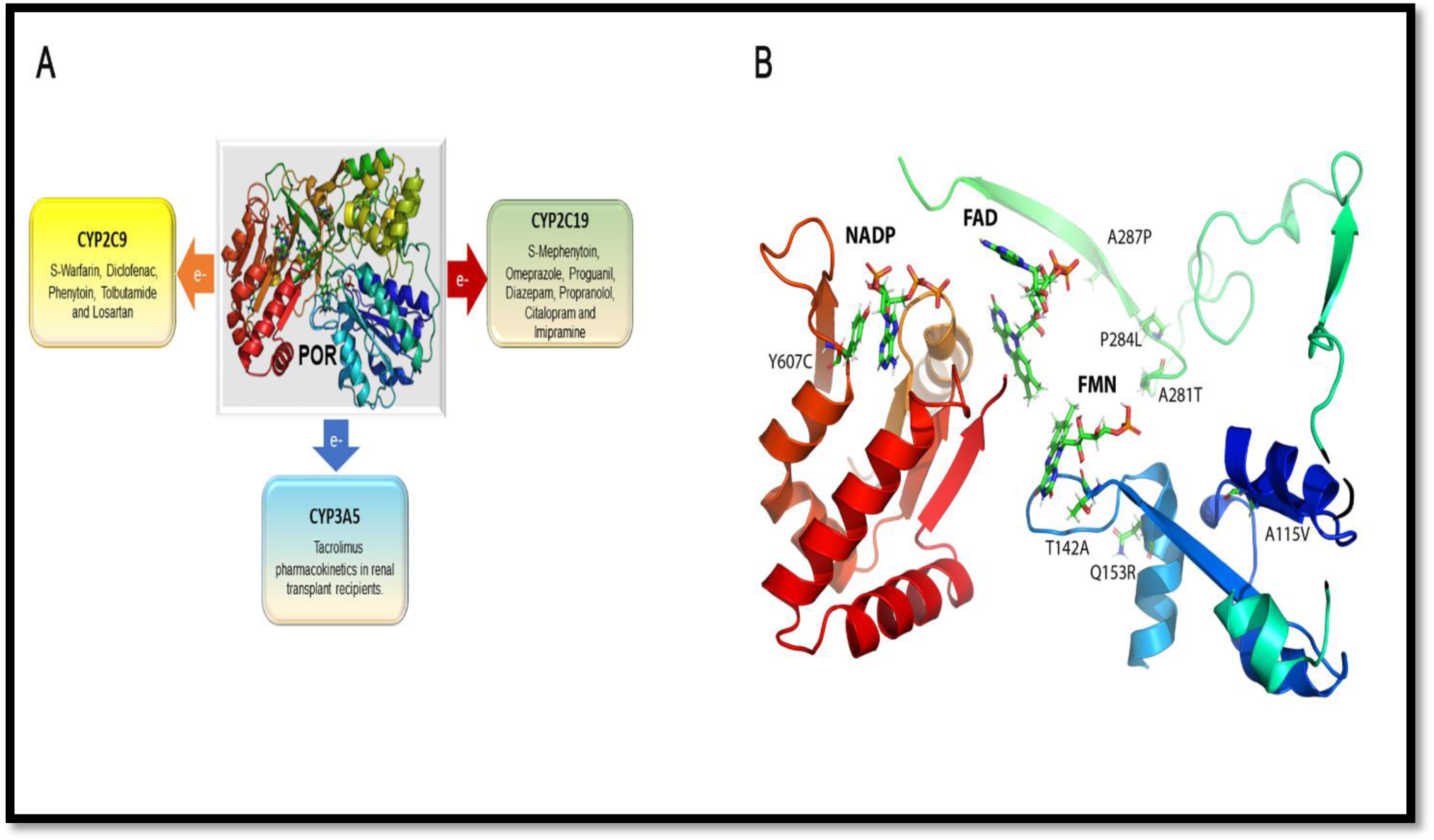
POR and location of variants studied. **A.** Role of POR in the metabolism of drugs by cytochromes P450 CYP2C9, CYP2C19, CYP3A5. POR supplies redox equivalents from NADPH to its redox partners via FAD and FMN co-factors. POR is the redox partner for cytochrome P450 proteins located in the endoplasmic reticulum, including the drug metabolizing enzymes CYP2C9, CYP2C19 and CYP3A5. A variety of drugs are metabolized by the three cytochrome P450 proteins tested in this study. Of note is the metabolism of warfarin and diclofenac by CYP2C9, metabolism of omeprazole, imipramine, proguanil, and diazepam by CYP2C19 and metabolism of renal transplant drug tacrolimus by CYP3A5. **B:** Location of variants studied. The model based on the x-ray crystal structure of the human POR (**PDB#3QE2**) showing the location of mutations studied in this report. The co-factors FMN, FAD, and NADP are shown as sticks. The mutation T142A is located next to the FMN binding site of POR, and its mutation would affect FMN binding. The Q153 residue is located at the surface of POR and predicted to have a role in the interactions with the redox partners. The A281, P284 and A287 residues are located in the hinge region of the POR. The hinge region of provides conformational flexibility, and mutations in the hinge region would result in conformational changes that would alter the interaction of POR with its redox partners.

Cytochrome P450 oxidoreductase deficiency (PORD) is a rare form of congenital adrenal hyperplasia (CAH) [12,13]. PORD causes defects of adrenal steroidogenesis in addition to several other symptoms [14,15,16,17,18,19,20,21]. Likewise, many other alterations in enzymes activities from mutations in POR has been observed due to the multiplicity of its redox partners, including heme oxygenase, CYP17A1, CYP19A1, and CYP3A4 [13,22,23,24,25,26,27,28,29]. Sequencing and biochemical studies have revealed many diseases caused by mutations in the POR gene [27,30,31,32]. These findings highlight the importance of POR variant studies on the enzymatic activity of different POR redox partners, especially the drug metabolizing enzymes. The cytochrome P450 CYP2C9 metabolizes approximately 15% of clinically relevant drugs, including warfarin, diclofenac, and losartan [33,34]. CYP2C19 is involved in approximately 2% of the clinical drug metabolism, such as anticonvulsant drugs like certain barbiturates [35,36,37]. Cytochrome P450 3A5 belongs to the subfamily 3 of human cytochrome P450 proteins, and along with CYP3A4, it is one of the most promiscuous enzymes. The CYP3A5 has a role in extrahepatic tissues, especially in intestine and gut [38,39]. CYP3A5 expression varies between ethnic groups due to an allele that leads to an incorrectly spliced mRNA and premature termination of translation. CYP3A5 is involved in tacrolimus pharmacokinetics and influences the rate of tacrolimus clearance, which is of importance in organ transplant [40].

We have been studying the variations in human POR genes since its link to disorders of steroid biosynthesis was discovered in 2004 [15,16,17,22,23,24, 25, 26, 27, 41, 42, 43]. These studies provided the background for exploring the effects of POR variations on different drug-metabolizing cytochrome P450 enzymes. In this study, we have investigated the effects on the activity of three important drug metabolizing cytochrome P450s, CYP2C9, CYP2C19 and CYP3A5 by potentially disease-causing variations of the POR gene identified from genome sequencing projects [12,13,32].

## Materials and Methods

### Recombinant expression of POR and Membrane purification

The human POR WT and variant forms of proteins (NCBI# NP_000932, Uniprot# P16435) were recombinantly expressed in bacteria using the previously described methods and expression constructs [24,26,27,44]. The protocol for expression of an N-27 form of POR variants and subsequent membrane purification have been described and adopted from our previous publications [23,26,27,28,29].

### Western blot analysis of POR content in the bacterial membranes

For Western blots, 1 μg of POR-WT and mutant membrane proteins were separated on an SDS-PAGE gel and blotted on to polyvinyl difluoride (PVDF) membranes to probe with a rabbit polyclonal antibody against wild-type human POR from Genscript (Genscript, NJ, USA) as described previously [43]. In all experiments described in this report, the normalized amount of POR content was used for the mutants as well as WT POR protein.

### Assay of cytochrome P450 CYP2C9 activity in reconstituted liposomes

The activity of CYP2C9 promoted by WT or mutant POR was tested using the fluorogenic substrate BOMCC (Invitrogen Corp, Carlsbad, CA, United States). The purified CYP2C9 (CYPEX, Dundee, Scotland, United Kingdom) was used to test the activities of the POR variants using 20 µM BOMCC as substrate. In vitro CYP2C9 assays were performed using a reconstituted liposome system consisting of WT/mutant POR, CYP2C9 and cytochrome b_5_ at a ratio of 5:1:1. The final assay mixture consisted of 5 µg DLPC (1,2-Dilauroyl-sn-glycero-3-phosphocholine) and proteins (1 µM POR: 200 nM CYP2C9: 200 nM b_5_), 3 mM MgCl_2_, 20 µM BOMCC in 100 mM Tris-HCl buffer pH 7.4 and the reaction volume was 100 µL. The P450 reaction was started by addition of NADPH to a final concentration of 1 mM, and fluorescence was measured on a Spectramax M2e plate reader (Molecular Devices, Sunnyvale, CA, United States) at an excitation wavelength of 415 nm and an emission wavelength of 460 nm for BOMCC.

### Assay of cytochrome P450 CYP2C19 activity in reconstituted liposomes

The activity of CYP2C19 promoted by WT or mutant POR was tested using the fluorogenic substrate EOMCC (Invitrogen Corp, Carlsbad, CA, United States). The purified CYP2C19 (CYPEX, Dundee, Scotland, United Kingdom) was used to test the activities of the POR variants using 20 µM EOMCC as substrate. In vitro CYP2C19 assays were performed using a reconstituted liposome system consisting of WT/mutant POR, CYP2C9 and cytochrome b_5_ at a ratio of 5:1:1. The final assay mixture consisted of 2.5 µg DLPC (1,2-Dilauroyl-sn-glycero-3-phosphocholine) and proteins (0.5 µM POR: 100 nM CYP2C9: 100 nM b_5_), 3 mM MgCl_2_, 20 µM EOMCC in 100 mM Tris-HCl buffer pH 7.4 and the reaction volume was 100 µL. The P450 reaction was started by addition of NADPH to a final concentration of 0.5 mM, and fluorescence was measured on a Spectramax M2e plate reader (Molecular Devices, Sunnyvale, CA, United States).

### Assay of cytochrome P450 CYP3A5 activity in reconstituted liposomes

The activity of CYP3A5 promoted by WT or mutant POR was tested using the fluorogenic substrate BOMCC (Invitrogen Corp, Carlsbad, CA, United States). The purified CYP3A5 (CYPEX, Dundee, Scotland, United Kingdom) was used to test the activities of the POR variants using 20 µM BOMCC as substrate. In vitro CYP3A5 assays were performed using a reconstituted liposome system consisting of WT/mutant POR, CYP3A5 and cytochrome b_5_ at a ratio of 5:1:1. The final assay mixture consisted of 5 µg DLPC (1,2-Dilauroyl-sn-glycero-3-phosphocholine) and proteins (1 µM POR: 200 nM CYP2C9: 200 nM b_5_), 3 mM MgCl_2_, 20 µM BOMCC in 100 mM Tris-HCl buffer pH 7.4 and the reaction volume was 100 µL. The CYP3A5 reaction was started by addition of NADPH to 1 mM final concentration, and fluorescence was measured on a Spectramax M2e plate reader (Molecular Devices, Sunnyvale, CA, United States) at an excitation wavelength of 415 nm and an emission wavelength of 460 nm for BOMCC.

### Protein structure analysis of POR variants

Three-dimensional structural models of POR (NCBI# NP_000932) proteins were obtained from the protein structure database (www.rcsb.org). We used the structures of the human POR (PDB # 3QE2) to analyze the location of amino acids described in this report [8]. Structure models were drawn using the software Pymol (www.pymol.org), and rendering of images was performed with POVRAY (www.povray.org).

### Statistical Analysis of results

Data are shown as mean, standard errors of the mean (SEM) in each group of replicates. Differences within the subsets of experiments were calculated using Student’s t-test. P values less than 0.05 were considered statistically significant.

## Results

### Sequence conservation and genetic distribution of POR variants

The POR variants studied in this report have been identified by mining the genome databases and performing structural stability and sequence conservation analysis to find out which of the variants may be potentially disease-causing [13]. Human POR is a 680 amino acid protein which has evolved from ferredoxin and ferredoxin reductase-like domains to form a single redox protein in eukaryotes. All variations in POR studied here are highly conserved across species and are found as rare variants in NCBI Exome database (**Table 1**). The A115V was found mainly in European samples while T142 is a rare variant identified in the American population. The P284L is also a rare variant of predominantly European origin while the A287P is the most common pathogenic variant of POR seen in patients of European origin. The Y607C is rare variant seen in European and Asian populations.

**Table 1:** Frequencies of POR variants from NCBI Exome database.

| POR | DBSNP id | Global | European | Asian | American | African |
| --- | --- | --- | --- | --- | --- | --- |
| **A115V** | rs199634961 | T=0.00020 | T=0.00028 | T=0.0002 | T=0.0000 | T=0.0001 |
| **T142A** | rs1304915832 | G=0.00000 | G=0.00001 | G=0.0000 | G=0.0001 | G=0.0002 |
| **Q153R** | - | - | - | - | - | - |
| **A281T** | - | - | - | - | - | - |
| **P284L** | rs72557938 | T=0.00005 | T=0.00004 | T=0.0001 | T=0.0000 | T=0.0000 |
| **A287P** | rs121912974 | C=0.00024 | C=0.00040 | C=0.0000 | C=0.0000 | C=0.0001 |
| **Y607C** | rs72557954 | G=0.00011 | G=0.00008 | G=0.0003 | G=0.0000 | G=0.0000 |

### Structural analysis of POR variants

The POR variants A115V, T142A, and Q153R, are located in the FMN binding domain of POR (**Figure 1B**). The POR variant A115V is predicted to be a structural variant which may alter redox partner interactions. The mutation T142A is located next to the FMN binding site of POR, and its mutation would affect FMN binding (**Figure 1B**). The Q153 residue is located at the surface of POR and is predicted to have a role in the interactions of POR with its redox partners. The A281, P284 and A287 residues are located in the hinge region of the POR. The hinge region in POR provides conformational flexibility, and mutations in the hinge region would result in conformational changes that may alter the interaction of POR with its redox partners.

### Effect on CYP2C9 enzyme activity by POR genetic variations

The A115V, T142A, and A287P variants of POR showed 6.5–6.8 % CYP2C9 activity compared to WT (**Figure 2**). POR variants P284L had only 22% of WT activity while the A281T and Y607C variants showed 27% of WT activity. POR variant Q153R showed 56% of WT activity in CYP2C9 assays.

**Figure 2:**
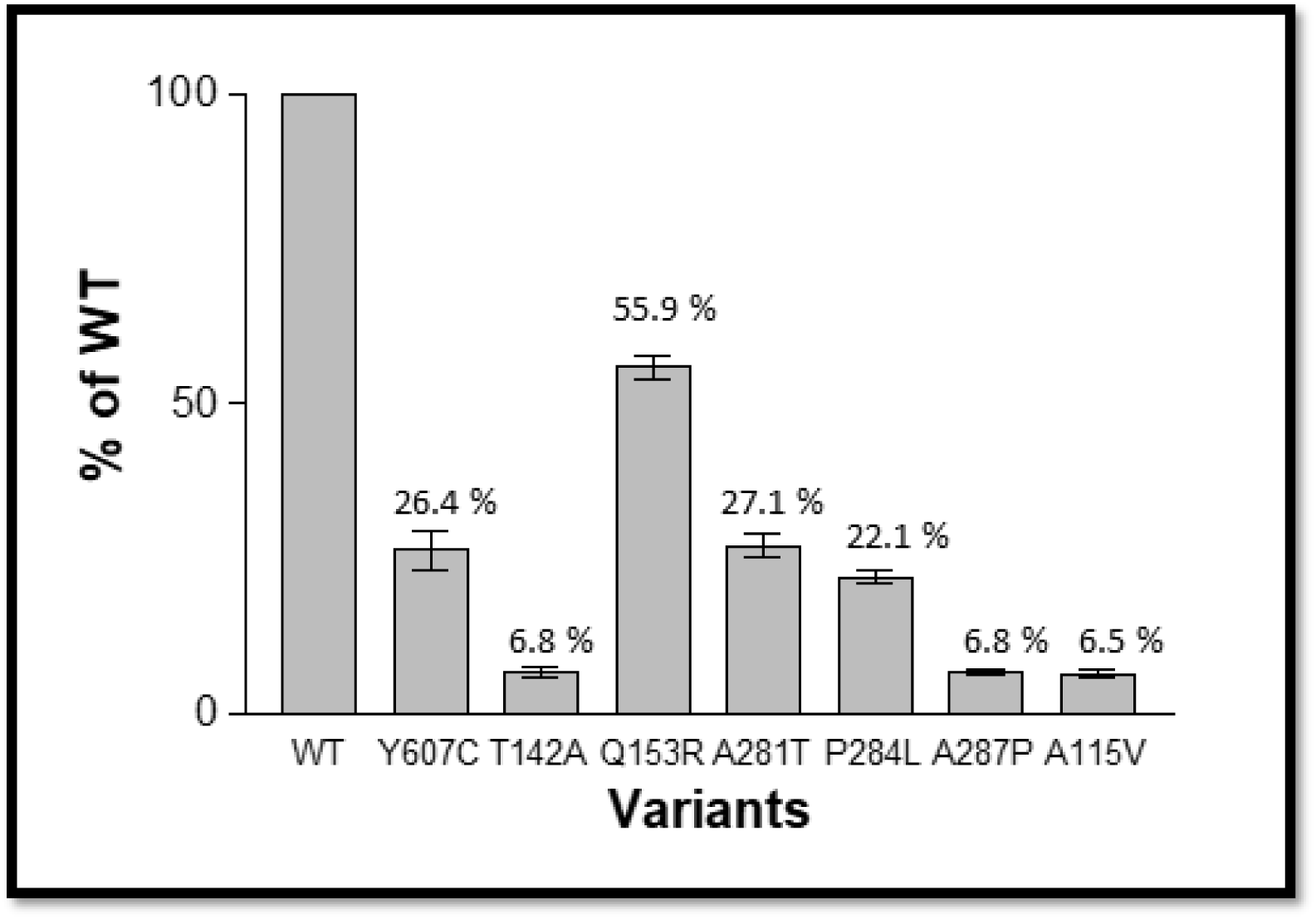
CYP2C9 activity promoted by WT and variants of POR proteins. Activity with the WT POR was set at a hundred percent, and results are given as a percentage of the activity supported by WT POR. In CYP2C9 assays POR variants A115V, T142A, and A287P had less than 10% of WT activity. POR variants A281T, P284L, and Y607C, lost around 75% of activity in CYP2C9 assays compared to WT POR. The POR variant Q153R retained 50% of WT activity. Data are presented as mean ± SEM of three independent replicates. P values obtained by students T-test were as follows: A115V 8.08E-11; T142A 2.44E-10; Q153R 8.12E-09; A281T 1.43E-07; P284L 1.06E-08; A287P 1.82E-11 and Y607C 1.21E-06.

### Effect on CYP2C19 enzyme activity by POR genetic variations

The A115V, T142A, A281T, P284L A287P, and Y607C variants of POR showed 41.5 –61.5 % CYP2C9 activity (A115V 43%, T142A 42%, A281T 62%, P284L 45% and A287P 42%) compared to WT (**Figure 3**). POR variant Q153R showed only a slight reduction in activity to 83% of WT POR (**Figure 3**). All variants had in general higher activities in CYP2C19 assays compared to CYP2C9 assays.

**Figure 3:**
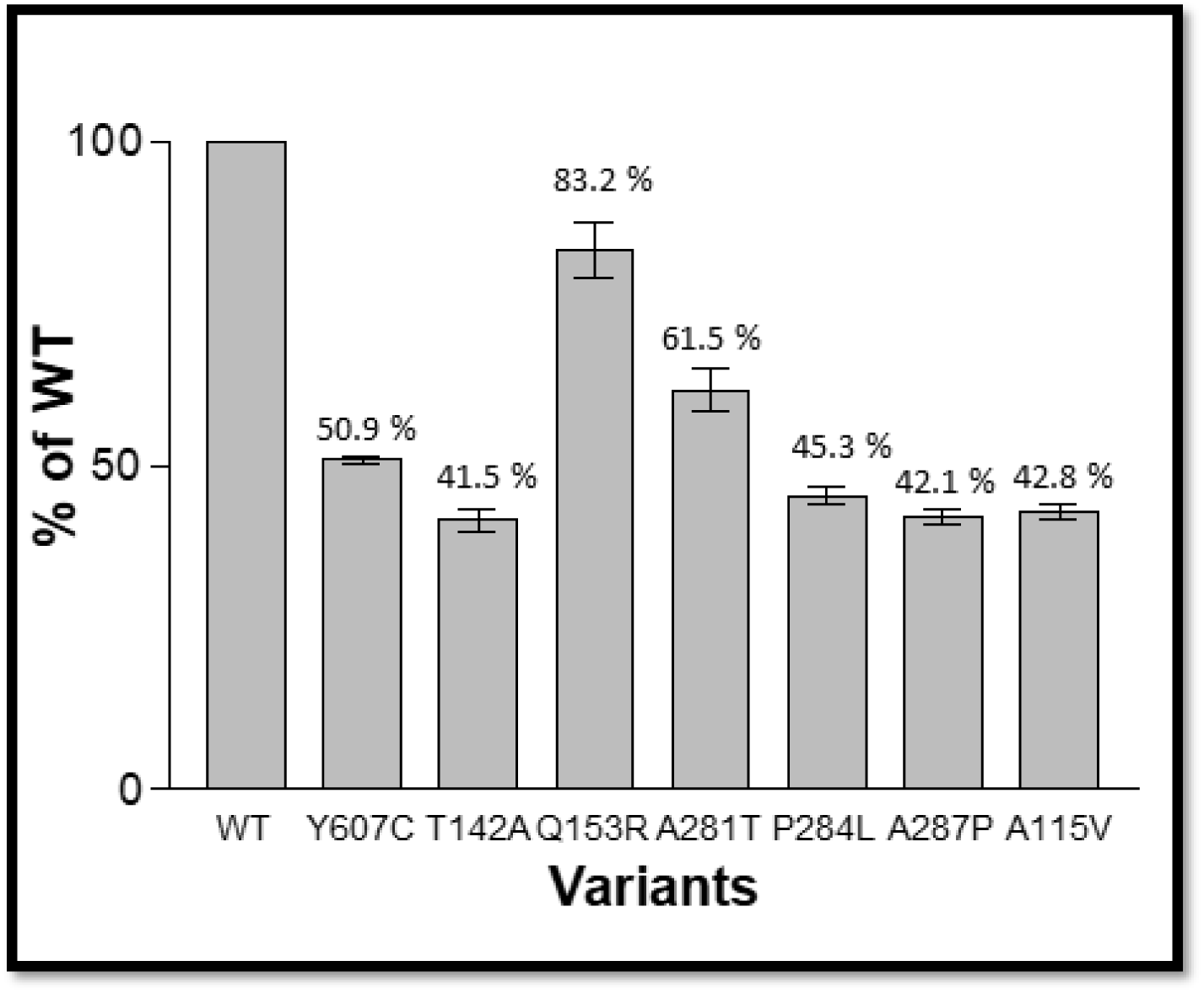
CYP2C19 activity promoted by WT and variants of POR proteins. Activity with the WT POR was set as a hundred percent, and results are shown as a percentage of WT activity. POR variants A115V, T142A, P284L, and A287P, showed more than 50 loss of CYP2C19 activity compared to WT POR. The variant Y607C had 51% of WT activity, and A281T variant had 62% of WT activity. POR variant Q153R showed 83% of WT activity in CYP2C19 assays. Data are presented as mean ± SEM of three independent replicates. P values obtained by students T-test were as follows: A115V 3.7E-08; T142A 2.1E-07; Q153R 0.005; A281T 4.1E-05; P284L 1.05E-07; A287P 3.4E-08 and Y607C 3.01E-09.

### Effect on CYP3A5 enzyme activity by POR genetic variations

The A115V, T142A, and A287P and variants of POR showed 15.8–23.1 % CYP3A5 activity (A115V 18%, T142A 16%, A287P 23%) compared to WT (**Figure 4**). The P284L had 44% of WT activity while the Y607C variant retained 67% of WT POR activity in CYP3A5 assays (**Figure 4**). POR variant A281T had moderately increased values compared to WT POR in CYP3A5 assays (138 %) but the Q153R variant of POR enhanced the CYP3A5 activity around 4 folds (**Figure 4**).

**Figure 4:**
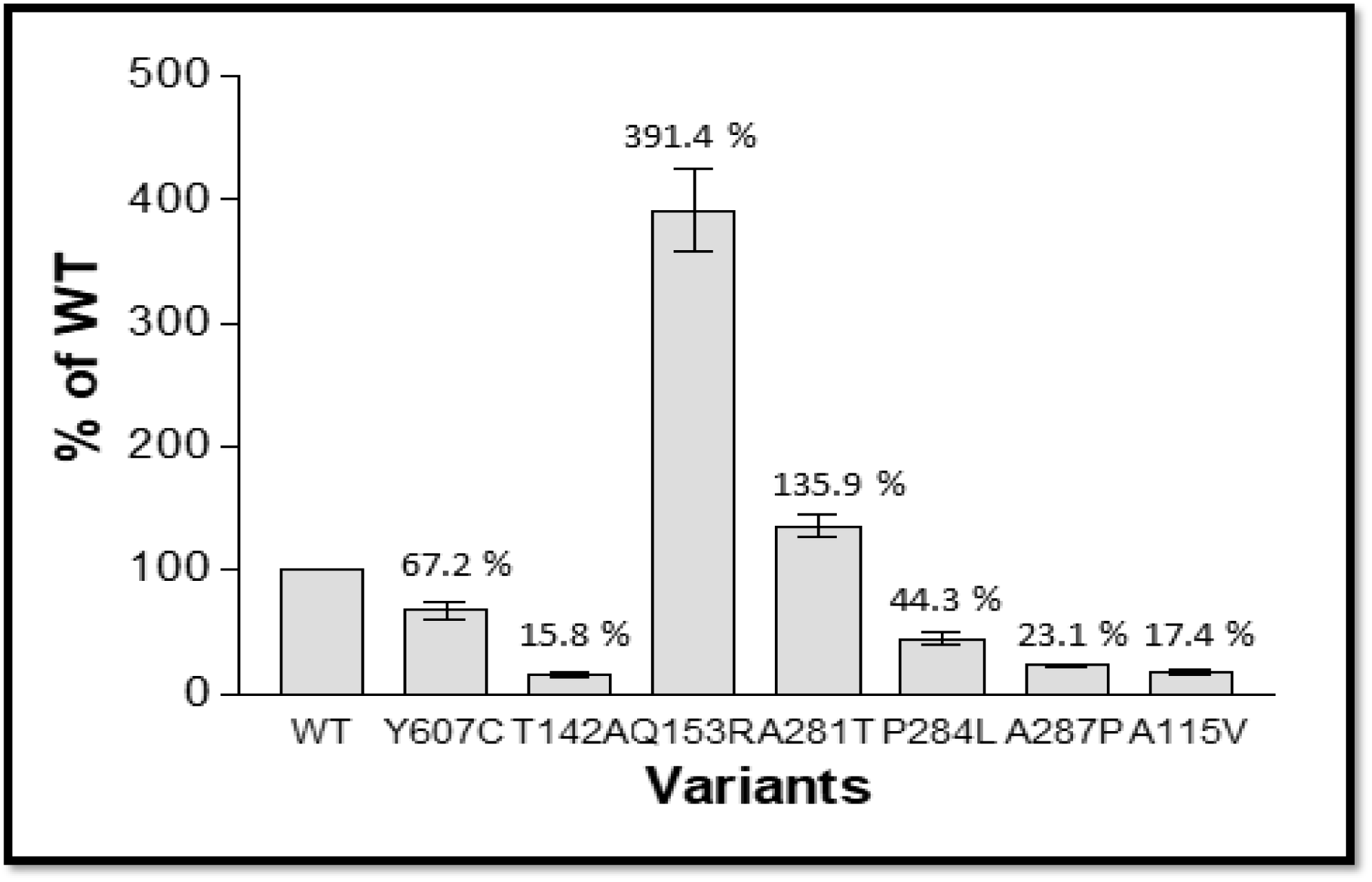
CYP3A5 activity promoted by WT and variants of POR proteins. Activity with the WT POR was set as a hundred percent, and results are shown as a percentage of WT activity. In CYP3A5 assays POR variant A115V had 17% of WT activity while variant T142A had 16% of WT activity. POR variant A287P had 23% of WT activity and variant Y607C had 67% of WT activity. Two variants showed higher than WT activity with A281T showing 136% of WT activity and Q153R showed 391% of WT activity in CYP3A5 assays. Data are presented as mean ± SEM of three independent replicates. P values obtained by students T-test were as follows: A115V 9.9E-13; T142A 1.28E-08; Q153R 1.24E-05; A281T 0.0024; P284L 7.26E-05; A287P 2.25E-08 and Y607C 0.0024.

## Discussion

Here we studied the effects on drug metabolism and toxicology caused by POR gene mutations identified as potentially disease-causing variations. We selected the POR gene variants based on the sequencing data available from the 1000 Genome Project and other large sequencing studies [13]. Our earlier studies on POR variants indicate that there may be considerable differences in the effects of individual POR variants on the activities of different redox partners. While most POR variants have been studied for effects on steroid metabolizing enzymes like CYP17A1 and CYP19A1, there are few studies that have examined the role of POR on drug metabolizing enzymes. We determined the enzymatic activities of three drug and xenobiotic metabolizing cytochromes P450 promoted by WT and the population genetic variants of POR. POR variant A115V earlier suspected as a polymorphism [27,32,42] inhibited all three cytochrome P450 activities severely, in agreement with its effect on CYP19A1 [43]. The A287P variant, which is the most common POR mutation found in patients of European origin [12,13,27,45], showed severe loss of activities in CYP2C9 and CYP3A5 assays which has important consequences for monitoring and treatment of patients [34,40].

Interestingly, POR variant Q153R showed a reduction of 20-50 activities with CYP2C9 and CYP2C19 but had a 400% increased activity with CYP3A5. The severe effects of A115V, T142A, and can be explained due to their proximity to the FMN binding site of POR that has a vital role in protein-protein interaction. The POR variant Q153R is located at the surface and considering its variable effects on different partner proteins; it is likely to have a role in the interaction with redox partners of POR. POR Variants P284L and Y607C inhibited activities of all P450 proteins tested. The A281T POR variant had less activity than the WT with CYP2C9 and CYP2C19 but showed slightly increased activity with CYP3A5. These results showed that POR variants have a diversity of possible effects depending on their redox partners.

These results indicate that detailed knowledge about the effects of POR genetic variations is necessary for correct diagnosis and treatment options for persons with POR deficiency and the role of POR in the alterations in drug metabolism needs to be addressed. Changes in drug and steroid metabolism due to genetic variations can be addressed using personalized metabolic profiling and supplementation using modified dosages of drugs, steroids, or co-factors [14,24,41,46]. Several drugs, including the organ transplant drug tacrolimus, have narrow therapeutic ranges, and severe loss of CYP3A5 activity due to mutations in POR requires considering POR variations as modifiers of drug metabolism in addition to cytochrome P450 variant alleles.

## Acknowledgments

This work was supported by grants from the Swiss National Science Foundation (31003A-135926), the Novartis Foundation for Medical-Biological Research (18A053) and Burgergemiende Bern to AVP. MNRV was supported by Consejo Nacional de Ciencia y Tecnología, Paraguay. We thank Prof. Walter L. Miller (UCSF, San Francisco, CA, USA) for providing several POR cDNA clones.

